# Impaired trap closure in the counting-deficient Venus flytrap mutant DYSCALCULIA is caused by cell wall biomechanics

**DOI:** 10.1101/2025.06.26.661685

**Authors:** Sonja Trebing, Matthias Freund, Anda Iosip, Célian Diblasi, Vincent Krennerich, Jonathan Kirshner, Mitsuhiko P. Sato, André Marques, Marie Saitou, Dirk Becker, Victor A. Albert, Ingrid Tessmer, Kenji Fukushima, Rainer Hedrich, Ines Kreuzer

## Abstract

Living in nutrient-poor environments, the carnivorous Venus flytrap *Dionaea muscipula* captures animal prey to compensate for this deficiency. Stimulation of trigger hairs located on the inner trap surface elicits an action potential (AP). While two consecutive APs result in fast trap closure in wildtype (WT) plants, sustained AP generation by the insect struggling to escape the trap leads to jasmonic acid (JA) biosynthesis, formation of the digestive “stomach”, and release of enzymes needed to decompose the victim. The *Dionaea muscipula* DYSCALCULIA (DYSC) mutant is able to fire touch-induced APs, but unlike WT plants, it does not snap-close its traps after two consecutive APs. Moreover, DYSC plants fail to properly initiate the JA pathway in response to mechanostimulation and even wounding, a well-known JA-dependent process conserved among plants. As demonstrated in previous studies, this DYSC mutant defect is associated with impaired decoding of mechanostimulation (i.e. touch) -induced Ca^2+^ signals. External JA application to the trap, however, restores slow trap closure and digestive gland function in DYSC, while rapid trap closure is JA-independent and cannot be rescued by exogenous JA application. Higher frequency mechanostimulation and thus more APs, however, revealed that DYSC is still able to close its traps, albeit much slower than WT plants. To reveal the molecular underpinnings of DYSC’s delayed trap movement, we generated a chromosome-scale *Dionaea* genome assembly and profiled gene expression. The refined transcriptomic analysis uncovered widespread misregulation of cell wall-related genes in DYSC, implicating altered cell wall plasticity in the sluggish mutant. Cell indentation studies by atomic force microscopy revealed a strictly localized and strikingly enhanced stiffening of the cell wall for DYSC that may hinder rapid trap closure and snap buckling. Together, these genomic, transcriptomic, and biophysical data identify cell wall elasticity as a key constraint on voltage and Ca^2+^ dependent trap kinetics. This finding documents the interrelationship between mechanosensing and Ca^2+^ signaling in the ultrafast capture organ of the Venus flytrap.

## Introduction

The carnivorous Venus flytrap, *Dionaea muscipula,* thrives in nutrient-poor environments where it captures animal prey to compensate for this deficiency. *Dionaea*’s exceptional hunting skills are based on fast electrical signaling and active trap movement. Displacement of one of the six trigger hairs located on the inner trap surface elicits an action potential (AP) which travels all along the trap^1^. The ion channels, pumps and carriers underlying *Dionaea*’s action potential have recently been identified and were linked to the different AP phases by molecular modelling and inhibitor studies^2–5^. The AP is accompanied by a simultaneous Ca^2+^-wave that is initiated by Ca^2+^ influx likely via GLR-type Ca^2+^ channels and by an ER-based calcium-induced calcium release (CICR)^4,6^. One single AP, i.e. only one trigger hair displacement (potentially triggered by a non-prey object), solely initiates a subcritical rise in cytosolic calcium concentration ([Ca^2+^]cyt). However, an actual prey animal visiting the trap provokes a second AP and thus a subsequent Ca^2+^ wave within 20-30 sec, while the first wave has not yet fully faded. Therefore, the rise in [Ca^2+^]cyt finally exceeds the threshold, triggering fast snap closure of the trap. In this manner, *Dionaea* avoids energetically costly trap movement in the absence of prey.

Captured prey will not be able to escape from the trap they cannot pass the barrier established by the entwined spines at the trap margin. Struggling to escape, a prey insect will repeatedly touch trigger hairs and thus evoke further APs and Ca^2+^ waves. Exceeding a second threshold of 3-5 APs, Ca^2+^ signaling translates into the activation of the jasmonic acid (JA) pathway: a rise in JA-Ile leads to stomach formation, i.e. tight trap closure and secretion of digestive fluids and transport proteins from gland complexes, permitting prey decomposition and prey-derived nutrient uptake^7^.

Rapid trap closure lasts only a fraction of a second and is brought about by hydraulically driven trap lobe deformation: the open, outward curved, concave trap lobes rapidly switch into a convex, closed conformation^8,9^. As prerequisite of this movement, the open trap is considered pre-stressed due to differences in turgor pressure between the inner and outer trap layers^9–11^. As a result, hydroelastic curvature energy is stored in the open trap. Rapid trap closure, also referred to as snap-buckling, is very likely brought about by i) slight shrinkage of the inner trap layer and ii) expansion of the outer trap layer, presumably facilitated by flow of water from inner to outer trap layers^9^. Trap closure can be inhibited by the application of aquaporin blockers^11^, as well as by dehydration of the plant^9^, thus pointing to the importance of turgor establishment and turgor changes during snap closure of *Dionaea traps*

Interestingly, a naturally occurring Venus flytrap mutant without snap-trapping function, initially called “ERROR”, was recently described^12^. This mutant – otherwise morphologically indistinguishable from wildtype plants – features traps that remain open when sensory trigger hairs are displaced two times. Because of its ‘counting’ defect leading to its impairment to snap close upon two touches, the mutant was renamed to ‘DYSCALCULIA’ (DYSC).

Remarkably, the APs evoked by stimulating DYSC mutant traps do not differ in shape, amplitude, and kinetics from those registered in wildtype plants. Therefore, the trap closure defect does not result from impaired hapto-electric signaling. Moreover, DYSC plants establish a WT-like Ca^2+^-signal in response to mechanostimulation. After mechanostimulation, however, the mutant fails to translate the counted APs and the associated Ca^2+^ signals into JA biosynthesis and thus does not enter the hunting cycle. Interestingly, DYSĆs impaired JA-response after mechanostimulation and wounding is restricted to the trap, while the petiole does not show this defect. External JA application to the trap, however, restores slow trap closure and digestive gland function. Transcriptomic analyses revealed differences in Ca^2+^-associated genes in WT and DYSC plants: upon trigger hair touch/AP stimulation, transcriptional activation of calcium signaling in DYSC traps was found largely suppressed. Therefore, DYSC mutants cannot properly read, count and decode touch-/AP-induced calcium signals, and thus they fail to properly translate touch-AP triggered Ca^2+^-signaling into the biosynthesis of JA ^12^.

Rapid trap closure via snap-buckling, however, is independent of JA, so that DYSĆs inability to snap shut its traps cannot be explained by the mutant’s incapability to properly initiate JA signaling in response to mechanoperception. Therefore, we asked if DYSĆs inability to snap close its trap might be based on morphological modifications affecting the trap’s ability to undergo the conformational changes required for snap buckling. Atomic force microscopy (AFM) based cell wall indentation studies revealed striking differences in cell wall elasticities between DYSC mutant and WT *Dionaea muscipula* traps. Transcriptomic analyses based on a newly generated chromosome scale *Dionaea* genome assembly confirmed that cell wall related genes are indeed massively deregulated in the DYSC mutant. DYSĆs inability to snap close its trap is therefore explained by cell wall biomechanical modifications affecting the trap’s ability to translate Ca^2+^-electrical signals in ultrafast snap buckling.

## Methods

### Plant materials

*Dionaea muscipula* (Droseraceae) WT plants were purchased from Cresco Carnivora (Netherlands). Before use, plants were acclimatized to greenhouse conditions with 20-22°C with a light:dark photoperiod of 16h:8h for a minimum of two weeks.

DYSCALCULIA (DYSC) mutant plants formerly known as “ERROR” cultivar, Rose and Basmati were purchased from Mathias Maier (https://green-jaws.com/). To overcome limitations in mutant plant material, axenic culture of DYSC was established by Dr. Traud Winkelmann (Leibniz University Hannover) and DYSC plants were further propagated inhouse in sterile medium (1/2 Murashige & Skoog^13^ without vitamins and MES, 2% Sucrose, 0.1% w/v 2-(N-morpholino)ethanesulfonic acid (MES), 1x Gamborg vitamins, pH 6.1, 0.6% Agar). Mature and healthy sterile plants were transferred to white peat and slowly acclimatized to greenhouse conditions over the course of multiple weeks. DYSC plants were grown under greenhouse conditions for at least one year before being used for RNA-Seq experiments.

### Mechanostimulation of WT & DYSC

APs were applied in different frequencies to WT / DYSC traps fitted with a white paper collar. Plants were left unstimulated for a minimum of 20 minutes before the start of the experiment to ensure an identical background for analysis of the hyperlapse videos. Special care was taken to ensure that no trigger hair (TH) was touched before starting mechanostimulation. Only one trap per plant was used. All plants were recorded with the same orientation for good visibility of the opening angle in a lightbox in front of a camera. Hyperlapse videos at 8x speed were recorded with Samsung Galaxy A71 camera. APs in DYSC traps were elicited by touching the trigger hairs with specific AP numbers and frequencies (1 AP/15 sec; 1 AP/30 sec; 1 AP/min; 10 or 20 APs in total). Trap closure was documented for at least 20 minutes for all samples; only for 20 AP with 1/min, videos were recorded for 30 minutes, so that each trap was left untouched for at least 10 minutes after mechanostimulation. Each sample was measured individually. While DYSC traps remained open during mechanostimulation and started closure with a delay, WT traps closed after 2 APs as expected. Therefore, WT Traps were only mechanostimulated twice with the specific frequency and no further APs were given after snap closure. The opening angle was determined every minute, and the data were analyzed manually in ImageJ (version 1.53f51). Sample sizes were n = 12 for DYSC and n = 6 for WT.

### COR Experiment

For coronatine (COR) treatment, 400 µl of 0.1 mM coronatine were sprayed on open WT and DYSC traps. Prior to spraying, each trap was fitted with a white paper collar and left for at least 20 minutes before the start of the experiment to ensure an identical background for analysis. Special care was taken that no TH was touched before spraying with COR. Only one trap per plant was used. All plants were recorded with the same orientation for good visibility of the opening angle in a lightbox in front of a camera. Trap closure was documented individually for 48 hours. Timelapse recording with 1 frame per minute was collected with a Canon camera (n = 10 per group). Opening angle measurement was conducted in ImageJ (version 1.53f51).

### EtOH Experiment

The Ethanol (EtOH) experiment was carried out in a light box to ensure constant and consistent lighting conditions. Experimental procedure was performed separately for WT and DYSC. Small glasses were filled with around 30 ml of 100% EtOH. Videos were taken with a Fujifilm x-T2 camera combined with Fujinon Aspherical lens Super EBC XF 18-55 mm. Videos of max. 30 minutes were taken per sample group. In total, 6 groups with 6 samples each were analyzed per plant type (n = 36). In order to observe different trap closure kinetics, traps had to be open. Therefore, whole leaves were cut at the lowest point of the petiole, as WT traps would have closed if trap and petiole had been separated. Immediately after cutting, the trap (incl. petiole) was placed in the ethanol-filled glass vessel. Only WT and DYSC traps with comparable opening angles and size were used. No trigger hair was touched during the process of cutting and placing into the ethanol solution. It was also ensured that the trap would go into the liquid directly and not come into contact with any external elements. Closure time was defined from first movement till closure without subsequent shape changes.

### Opening Angle and lobe curvature Measurements

Opening angles before and after trigger hair stimulation were quantified by timelapse videos recorded by a Samsung Galaxy A71 camera. While WT plant movements were recorded using standard speed videos, DYSC mutant hyperlapse videos were recorded at 8x speed. Videos were analyzed with ImageJ (version 1.53f51). The use of three reference points (left lobe rim (outer part), midrib (lower part) and right lobe rim (outer part)) ensured consistency between the analysis of the measurements. Angle measurements were taken for each sample at one-minute intervals. Curvature measurements of open trap lobes were performed with ImageJ Kappa-plugin. A curve was manually created to match the curvature of the trap lobes. The dimensions of both lobes of each trap were measured and subjected to analysis.

### AFM

For cell wall elasticity measurements, atomic force microscopy (AFM) was used. The preparation of the WT and DYSC traps was conducted in an identical manner. Each sample was prepared directly before measuring to ensure same time interval and minimal water loss between preparation and measurement. Traps were cut at the transition to the petiole. Once the trap had been separated into two trap lobes by a cut set along the midrib, it was necessary for steric reasons to remove the remaining components of the midrib to allow for placement of the individual lobes under the AFM mechanical probe. Individual lobes were attached to the center of a microscope slide using double-sided adhesive tape. Trap lobes were attached to the adhesive tape either by their inside or by their outside surfaces, depending on which side of the lobe was intended for elasticity measurements (the facing-up side). Three different positions were probed: directly below the rim, in the middle between the two trigger hairs, and near the former midrib. To ensure that differences in force-indentation behaviour were not due to prolonged time after cutting of the lobes, the order in which the different positions were measured was altered between samples. At each position, 10 force-indentation curves were collected in direct succession. Measurements were performed using a Molecular Force Probe 3D (MFP-3D) AFM (Asylum Research, Oxford Instruments) in contact mode and SD-R150-NCL AFM probes harboring a spherical tip with and end diameter of approximately 150 nm diameter (Nanosensors, nominal resonance frequency ∼ 190 kHz and spring constant ∼48 N/m). Force-indentation curves were obtained with a fixed force distance of 2 µm, trigger point of 10 nm, and tip approach/ retract velocity of 2.0 μm/sec. The force-indentation curves were analyzed with MFP software. From the recorded force curves Young’s Moduli were obtained via the Hertz model (Equation 1), where Ec is the overall measured Young’s modulus of the entire system and E1 and E2 are the contributing Young’s moduli of the sample and of the AFM tip cantilever, respectively. υ1/2 are the Poisson ratios of the sample and of the AFM tip material (Silicon, υ2 = 0.17). We used a Poisson ratio (υ1) of 0.3 for the cell wall based on previous reports for other plant cells^14^.

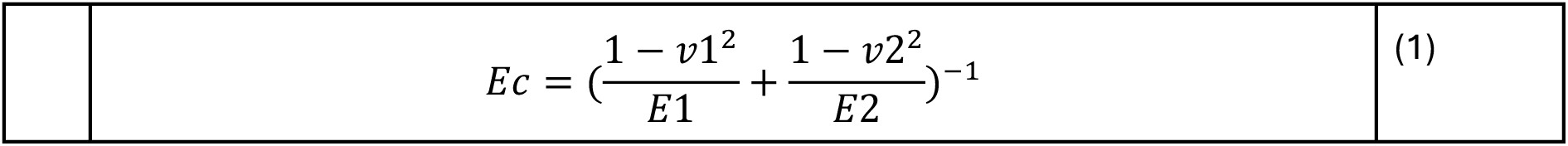

### Transcriptome sequencing

For RNA-Seq ground state comparison, untouched and healthy traps of WT and DYSC were used. Each group consisted of 3 replicates. RNA was isolated essentially as described before^5^. RNA-Seq was performed paired end (150 bp) by GATC Biotech (nowadays Eurofins Scientific) on an Illumina HighSeq2000. Quality check was done by fastQC (version 0.12.1) and multiQC (version 1.12)^15,16^. Paired end reads were mapped to the newest available *Dionaea* genome version (described below). The mapping procedure was conducted with AMALGKIT (version 0.12.0, https://github.com/kfuku52/amalgkit), utilizing the toolkit with default commands from ‘amalgkit integrate’ to ‘amalgkit merge’. The resulting table was used to work in RStudio (version 4.3.3) with DESeq2 package (version 1.4.4)^17^. Differential expression analysis is based on the comparison between DYSC (Condition A) versus WT control samples (Condition B) using DESeqDataSetFromMatrix function. Genes that passed the following filters were considered as differentially expressed genes (DEGs): for upregulated genes: padj < 0.05, log2FC > 1, base mean of Condition A > 50 counts and for downregulated genes: padj < 0.05, log2FC < −1, base mean of Condition B > 50 counts.

### MapMan bin annotation and enrichment analysis

Functional gene annotation was done with Mercator4 (version 6) and Mercator v3.6^18–21^. The *Dionaea* genome FASTA file was processed using the available online tools (https://www.plabipd.de/mercator_main.html) for Mercator v3.6 and Mercator4 v6, with the objective of conducting functional annotation. Furthermore, Mercator4 BIN enrichment analysis was completed via the tab located on the left-hand side of the website. For enrichment analysis Two-sided Fisher’s exact test was performed and an FDR adjusted p-value cutoff of 0.05 was set. In addition to the Mercator4 mapping file, the genes of interest were provided, and the background genes consisted of the entire genome.

### High-molecular-weight genomic DNA isolation

A stock solution of the nuclear isolation buffer (IB) was prepared as follows: 15 mM Tris-HCl, 10 mM EDTA-2Na, 130 mM KCl, 20 mM NaCl, 8% (w/v) PVP-10, 250 mg/L spermine tetrahydrochloride, and 350 mg/L spermidine trihydrochloride, with pH adjusted to 9.4. Then, 100 ml IB was mixed with 7.5 ml of β-mercaptoethanol and 100 μl of Triton X-100 to prepare the IBTB buffer. Fresh leaf tissues were frozen in liquid nitrogen and were ground into fine powders with a mortar and a pestle. An aliquot of 3 g tissue powders was then transferred into 30 ml of an ice-cold IBTB buffer in a 50-ml tube, and the tube was vigorously mixed until the solution was homogenized. The homogenate was gently mixed with a magnetic stir bar at 100 rpm for 10 min on ice. The homogenate was then strained through a nylon filter with the mesh size of 100 μm (Falcon® Cell Strainers, Corning) to remove tissue fragments. This step was repeated with a nylon filter with the mesh size of 40 μm. An aliquot of 100 μl of Triton X-100 was added per 10 ml of homogenate, and the tube was gently agitated by inversion until completely mixed. The tube was then centrifuged at 4°C at 2000 ×g for 10 minutes to pellet the nuclei. Supernatant was discarded. The nuclei pellet was then subjected to the DNA extraction using NucleoBond HMW DNA kit (Macherey-Nagel) according to the manufacturer’s instruction of the ‘Enzymatic lysis’ protocol. DNA was eluted with 150 μl of HE buffer provided by the kit. DNA was quantified with the Qubit dsDNA BR kit (ThermoFisher Scientific).

### Genome sequencing for assembly

Wild-type genomic DNA was sequenced with the Oxford Nanopore technology (ONT) by Novogene UK. Briefly, the HMW DNA fragments were end-polished, nick repaired and A-tailed. The 1D library itself was produced by ligating sequencing adapters onto double-stranded DNA fragments. After purification with AMPure XP beads, the prepared library was examined with Qubit 3.0 fluorometer (Life Technologies, USA) for quantification and BioAnalyzer for size distribution detection. The libraries were sequenced on the PromethION platform with eight PromethION flowcells (R9.4.1 FLO-PRO002).

### Genome assembly

A total of 694-Gb ONT reads (Data S1B) were assembled into contigs using Flye v2.8.3 ^22^ with the options --asm-coverage 100 and --genome-size 3.18g. While the flow-cytometric genome size estimate of 3.187 Gb was obtained from previous literature^23^, we determined a *k*-mer-based estimate of 2.55 Gb using GenomeScope v2.0 (https://github.com/schatzlab/genomescope)^24^. The genome assembly was then polished using medaka v1.4.3 (https://github.com/nanoporetech/medaka) with the option --model r103_prom_high_g360 and all ONT reads as input and subsequently using NextPolish v1.3.1 (https://github.com/Nextomics/NextPolish)^25^ with previously generated 294-Gb Illumina short reads (ERR3638806–ERR3638809). Repeat sequences were masked by RepeatModeler v2.0.2 (https://github.com/Dfam-consortium/RepeatModeler)^26^, and then allelic contigs were collapsed using Purge Haplotigs v1.1.2 (https://bitbucket.org/mroachawri/purge_haplotigs/src)^27^ with the ONT reads as input. The HiRise scaffolding with Omni-C libraries (Data S1B) was conducted by Dovetail Genomics (CA, USA). Misjoints were corrected manually. Scaffolds were taxonomically assigned using MMseqs2 v13.45111 (https://github.com/soedinglab/MMseqs2)^28^, and contaminated sequences were removed.

### Repeat masking

For subsequent gene model prediction, repetitive elements on the reference genome were masked using RepeatMasker v4.0.9 (https://github.com/Dfam-consortium/RepeatMasker^29^) with a species-specific repetitive sequence library generated by RepeatModeler v2.0.2 (https://github.com/Dfam-consortium/RepeatModeler)^26^.

### Gene model prediction

Gene model prediction was performed using a combination of *ab initio* and homology-based approaches using funannotate v1.8.14 (https://github.com/nextgenusfs/funannotate^30^). For RNA-Seq–guided gene model training, Illumina RNA-Seq reads (Data S1A) were provided as input to the ‘funannotate train’ pipeline. This step aligned RNA-Seq reads to the reference genome and performed *de novo* transcriptome assembly using Trinity v2.8.5^31^ to generate high-quality transcript evidence. The derived transcript models were used to refine gene predictions and inform splice site determination. Default parameters were generally used, with the maximum intron length (-- max_intronlen) set to 20 kb. Gene prediction was conducted using the ‘funannotate predict’ pipeline with the repeat-masked genome and transcript evidence. Parameters included the use of the “embryophyta” BUSCO dataset for gene model quality assessment and the optimization of *ab initio* gene prediction tools such as Augustus v3.3.3 (https://github.com/Gaius-Augustus/Augustus)^32^ and GeneMark-ES v4.71 (https://genemark.bme.gatech.edu/gmes_instructions.html)^33^. Evidence-based gene models were generated by integrating *ab initio* predictions with aligned transcripts and protein homology data. To further refine the gene models, funannotate update was applied, which leveraged the previously generated transcript and protein evidence to resolve potential structural inconsistencies, ensuring more biologically accurate gene models. Functional annotation of predicted coding sequences was performed with Trinotate v3.2.1 (https://github.com/Trinotate/Trinotate)^34^ with the DIAMOND BLASTP search (v2.1.9, E value cut-off = 0.01, https://github.com/bbuchfink/diamond)^35^ against the UniProt database downloaded on 21 June 2022.

### Mutant genome resequencing

Genomic DNA samples were obtained from *Dionaea* mutants as described above and were sequenced by Novogene UK (Data S1B). Briefly, genomic DNA were subjected to 350 bp insert DNA library preparation using the NEBNext® Ultra™ II DNA Library Prep Kit (Cat No. E7645) following the manufacturers’ instructions. The constructed libraries were purified with the AMPure XP system (Beckman Coulter, Beverly, USA), checked for size distribution using an Agilent 2100 Bioanalyzer (Agilent Technologies), and molarity was quantified by qPCR. Libraries were sequenced on an Illumina NovaSeq 6000 platform using S4 flow cells with PE150.

### Detection of single-nucleotide variations

For single nucleotide variations (SNV) detection, genomic and transcriptomic reads of each genotype were separately mapped to their reference genome using BWA-MEM^36^ (https://github.com/lh3/bwa). Genotype likelihoods were calculated with BCFtools v1.15.1^37^ (https://github.com/samtools/bcftools), followed by variant calling using the same software version. SNVs were subsequently annotated and filtered^38^ (https://github.com/pcingola/SnpEff), applying criteria that included a minimum quality score of 30, homozygosity, and a depth between 3 and 2× the average read coverage. Annotated SNVs (Data S1F) were required to satisfy criteria for both genomic and transcriptomic datasets and to be unique to the individual.

### Detection of structural variants

Structural variants (SVs) were identified independently for each individual using Manta v1.6.^39^ (https://github.com/Illumina/manta). The resulting SVs were subsequently merged across all individuals and genotyped using BayesTyper v1.5^40^ (https://github.com/bioinformatics-centre/BayesTyper), treating all contigs as diploid and without specifying any sex chromosomes. Variants with a quality score below 10 were discarded, as were variants classified as “Unknown” across all three individuals. This filtering was performed using VCFtools v0.1.16^41^ (https://github.com/vcftools/vcftools).

### Gene Ontology (GO) functional enrichment analysis

Gene models from *Dionaea* were annotated with the highest alignment score matches using BLASTP versus *Arabidopsis* protein sequences, using an *e*-value cutoff of 1 × 10^−5^. For GO enrichment analyses, tandem duplicates from the *Dionaea* assembly were downloaded from CoGe’s SynMap tool ^42,43^, and those annotatable at the above threshold were used as foreground subsets for GO enrichment analysis with GOATOOLS ^44^, using all similarly annotatable genes in the genome as background. Results were Bonferroni-adjusted, and p < 0.05 was used as the threshold for significance.

## Results

### DYSC can close their traps albeit much slower than WT

During numerous mechanostimulation experiments, we observed that few DYSC mutant plants – even though they did not snap close their traps like wildtype plants – had slightly reduced opening angles between their two trap lobes upon trigger hair stimulation. Therefore, the DYSC mutants were able to initiate trap closure but could not snap close their traps like WT traps do. Since this feature is barely visible to the naked eye, we monitored the opening angle of wildtype and DYSC traps over time using time-lapse imaging. The opening angles of WT and DYSC plants were recorded over a period of 20 minutes after mechanostimulation at frequencies of 1 AP / 15 sec, 1 AP / 30 sec, or 1 AP / 60 sec (Figure 1 and Figure S1A, B). As expected, all WT traps snap closed after 2 APs. When eliciting 10 APs (Figure 1A, C and Figure S1A), DYSC trap movement was most pronounced at a frequency of 1 AP / 60 sec. Under these conditions at least 5 out of 12 DYSC traps managed to reduce their opening angle over time (Figure 1C). However, even in this group, many of the mutants were still completely open or showed only minimal changes in trap geometry. A rise in stimulation frequency to 1 AP / 30 sec or 1 AP / 15 sec did not enhance DYSC trap closure. Since the elicitation of 10 APs at 1 AP / 60 sec is our standard mechanostimulation setup ^12^, we checked if trap DYSC closure could be promoted by the application of more than 10 APs. In response to 20 APs, DYSC traps closed further than after 10 APs at all tested frequencies (Figure 1 and Figure S1). 20 trigger hair displacements at 1

**Figure 1.**
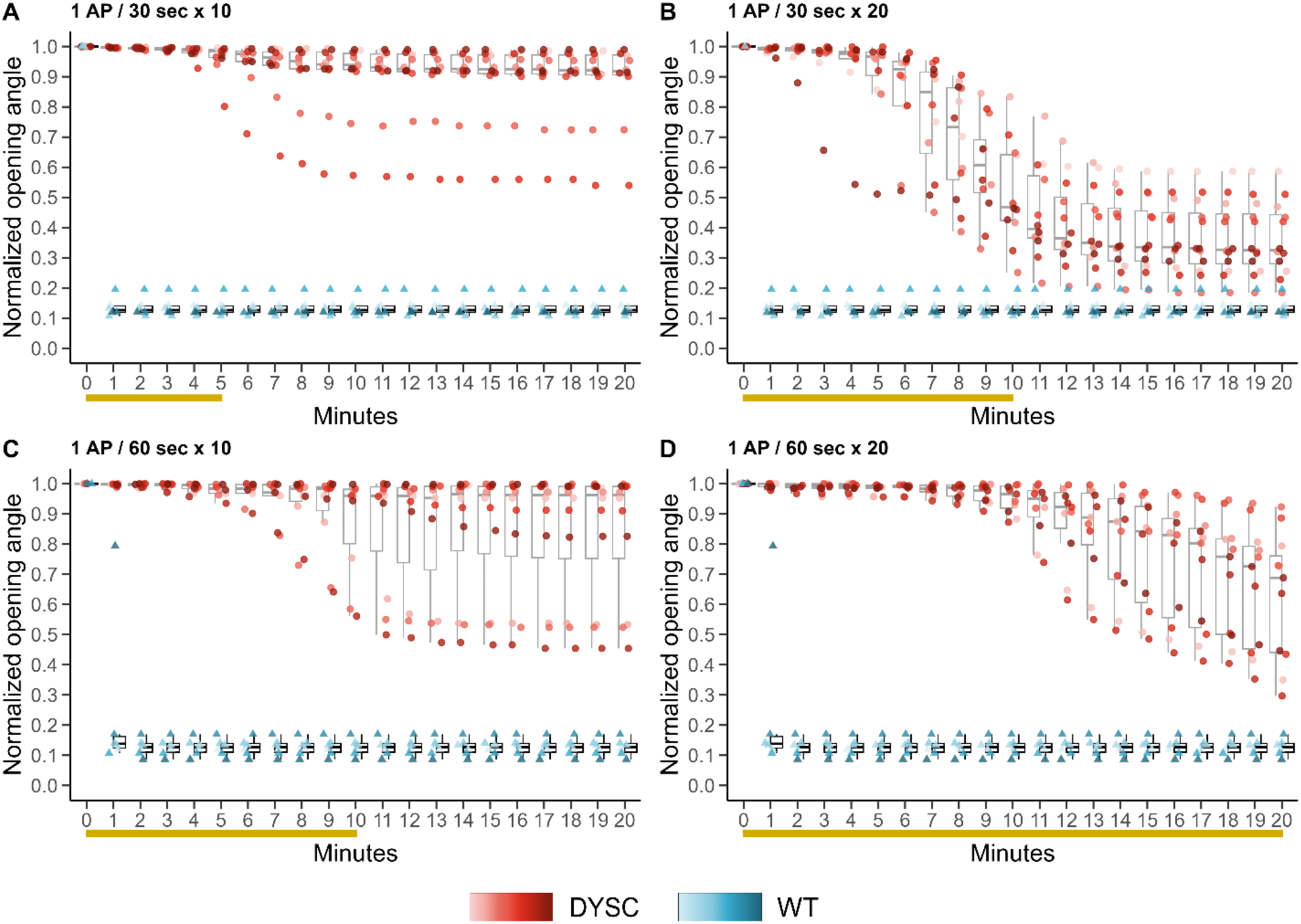
Changes in trap opening angles with different frequencies and total amount of AP for DYSC and WT. Normalized opening angle after mechanostimulation of traps with 10 AP or 20 AP with different frequencies for WT (blue) and DYSC (red). Left side (A, C): administration of a total of 10 APs to samples, right side (B, D): administration of a total of 20 APs. Each opening angle per sample at a given time was normalized to the maximum opening angle before starting mechanostimulation treatment. n = 12 for DYSC and n = 6 for WT. For normalization, 1 = open trap and 0 = sealed trap. 2 APs in WT samples led to fast trap closure in all cases. Orange bars below the x-axis indicate duration of mechanostimulation for DYSC samples with the specific frequency and AP number shown in the header of each graph. WT = *Dionaea muscipula* wildtype. DYSC = *Dionaea muscipula* DYSCALCULIA mutant/ cultivar. See also Figure S1.

/ 30 sec even resulted in detectable movement in all DYSC traps tested. While some of them were completely closed 15 min after mechanostimulation onset, others reduced their opening angle at least by 50% (Figure 1B). In response to 20 AP at 1 AP / 30 sec, DYSC traps had reached their maximum trap closure (i.e. minimum opening angle) 20 min after stimulation onset and thus 10 min after cessation of trigger hair bending. To test whether traps that had been stimulated with 20 APs at 1 AP / 60 sec would further close after mechanostimulation had been terminated, we extended the observation period for this specific sample group to a duration of 30 minutes (Figure S1E). However, the opening angles of these DYSC traps did not narrow further. Therefore, we concluded that DYSC trap closure can be induced by mechanostimulation in principle, although the mutant traps are much slower in doing so, and do not snap close. While WT plants close their traps by rapid conformational change of their traps, DYSC trap closure is not associated with WT-like snap buckling. Moreover, complete trap closure in DYSC more likely occurs at higher AP numbers applied at a higher frequency than is usually required for WT mechanostimulation. Finally, the closing process in DYSC seemed to require sustained trigger hair displacement and stopped when mechanostimulation was ceased.

### DYSC closure in response to JA stimulation is delayed

As described earlier, DYSC plants can be forced to close their traps by the application of the JA-mimic coronatine (COR), thus bypassing mechanostimulation induced by trigger hair bending, and instead entering directly into the secretion phase of its hunting cycle^12,45^. We therefore examined trap closure over time in WT and DYSC after spray application of 100 µM COR. As expected, COR treatment induced slow trap closure in both WT and DYSC (Figure 2). However, the majority of WT Traps were fully closed within the first 14 h following spray application, whereas this process took at least 20 h in most DYSC traps, again indicating slower trap closure in the mutant. Regarding curve progression, the (negative) slope of WT trap closure appeared much steeper than that in DYSC traps (Figure 2A). On average, WT traps closed at a speed approximately 2x faster than DYSC traps. While WT traps closed within 5 h, the same process took more than 10 h in DYSC plants (Figure 2B and C). Taken together, DYSC traps can close in response to COR treatment, but here again, the closure process takes much longer: i.e. their closing speed is significantly reduced compared to WT.

**Figure 2.**
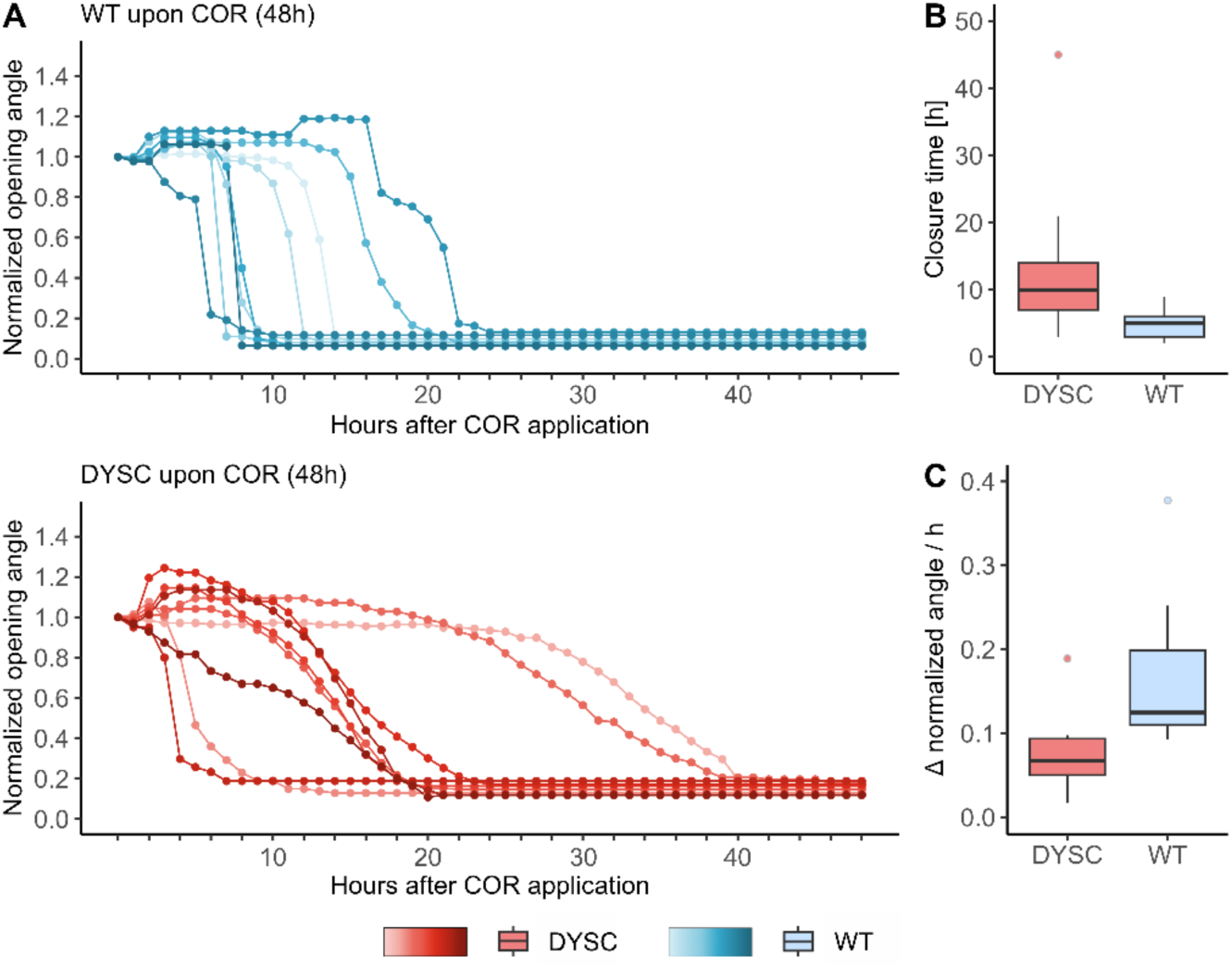
DYSC closure in response to COR application is effective but much slower than in WT traps. (A) Change in opening angle of traps sprayed with 100 µM coronatine (COR) in WT (blue) and DYSC (red) over 48h. The initial opening angle differed slightly between individual traps and was thus normalized to 1. (B) The closure time represents the slope in A; shown is the time between normalized opening angle < 0.9 and the minimum for each sample. (C) COR-induced closure speed in WT and DYSC. Boxplots include the median (horizontal line), the upper / lower quartiles and whiskers. Colored dots show the outliers per analyzed group. n = 10 for both WT and DYSC. WT = *Dionaea muscipula* wildtype. DYSC = *Dionaea muscipula* DYSCALCULIA mutant/ cultivar.

### Closure duration in DYSC is generally prolonged

*Dionaea* WT traps can be harvested by carefully cutting them from the plant with their petioles still attached to the capture organ. Under these conditions, the traps usually stay open, whereas they instantaneously close if the trap is directly cut off its petiole. However, we noticed that WT traps still attached to the petiole snap close immediately when submerged in 100% ethanol (EtOH). This treatment kills living cells by disrupting all membranes, thus allowing to analyze trap movement solely determined by cell wall properties. Therefore, both WT and DYSC leaves (traps still attached to their petioles) were subjected to submersion in EtOH and the duration required for the traps to close from initial movement to completion was monitored (Figure 3). While WT traps initiated closure 21 sec after immersion, the onset of closure was only slightly delayed in DYSC (24 sec). However, closure duration of WT traps was markedly shorter than that of the DYSC mutant: while most WT traps snap closed within 4 sec after initiation of closure, the closure process was significantly prolonged to 340 sec in DYSC traps. Therefore, DYSC plants may feature cell wall properties interfering with proper trap closure by snap buckling.

**Figure 3.**
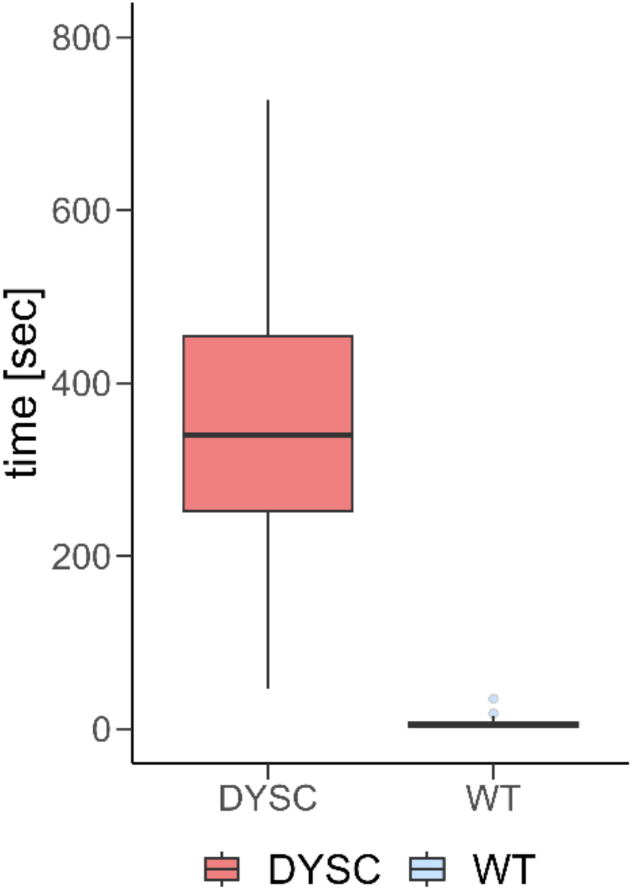
Submerging traps in 100% EtOH results in a longer closure duration for DYSC. Closure duration of *Dionaea muscipula* traps submerged in 100% EtOH. The boxplot shows closing time (sec) for WT (blue) and DYSC (red). Closure duration was defined as the interval from first movement of trap lobes until the spines at the trap margin crossed. Boxplots include the median (horizontal line), the upper / lower quartiles and whiskers. Colored dots show the outliers per analyzed group, n = 21-28 for WT / DYSC. WT = *Dionaea muscipula* wildtype. DYSC = *Dionaea muscipula* DYSCALCULIA mutant/ cultivar. See also Figure S2, Videos S1 and S2.

A condensed illustration of the experiment is presented in video S1. Upon closer examination of the recorded videos, we observed that DYSC traps exhibited less pronounced conformational changes compared to WT (Video S2). As described above, WT traps usually undergo a hydraulically driven lobe deformation, transferring the open, concave trap lobes into the convex, energetically favorable closed state. DYSC traps, however, are characterized by less curved traps and a smaller opening angle already in the open ground state (Figure S2), which could explain less pronounced changes in DYSC trap conformation during closure.

However, DYSC traps are impaired in snap closure which might be explained by altered cell wall properties compared to the WT. We therefore aimed to investigate the genomic modifications underlying this defect.

### Chromosome-scale genome assembly

As mentioned above, the DYSC mutation has been associated with defects in calcium decoding and Ca^2+^-regulated JA-biosynthesis in a previous study^12^. However, the *Dionaea* genome^23^ used as the reference in that work was highly incomplete, missing several essential genes, including the gene encoding the JA receptor *CORONATINE INSENSITIVE 1* (*COI1*). We therefore generated an all-new genome assembly using long-read DNA sequencing with Oxford Nanopore Technology, complemented by chromatin conformation capture sequencing with Omni-C technology.

This approach provided a genome assembly comprising 16 chromosome-scale scaffolds (Figure 4a-b), consistent with the reported chromosome number^46^, achieving a scaffold N50 of 144.5 Mb. The total assembly length of 2.55 Gb precisely matched the *k*-mer-based genome size estimate (2.55 Gb, Figure S4). Notably, the assembly recovered over 1 Gb of previously unsequenced genomic regions^23^, which were characterized by higher GC content (Figure 4c). Gene model prediction identified 38,887 genes, with a BUSCO completeness score of 84.3%, representing a 12.0% improvement over the previous assembly. Furthermore, this modelling included over 17,000 additional gene predictions compared to the earlier gene set^23^ (Figure 4c), with an increased proportion of functionally annotated genes (Figure 4d). These advancements suggest that prior characterizations of the *Dionaea* genome as gene-poor^23^ were likely artifacts of suboptimal assembly quality rather than a genuine paucity of genes. In sum, our new chromosome-scale assembly provides a robust and comprehensive gene set, facilitating more detailed analyses of carnivory-related traits and enabling (e.g. transcriptomic) investigations with enhanced genomic resolution. As an example, *Dionaea* COI1 is not only encoded within our new chromosome-scale assembly, but it is also represented by two tandemly duplicated genes, whereas the syntenic regions in the Arabidopsis and *Vitis vinifera* genomes contain only one *COI1* homolog each (Figure S3). Moreover, and also affirming to the utility of our assembly, the *COI1* gene pair was revealed by anonymous Gene Ontology (GO) enrichment analysis of tandem duplicates that were collaterally output from self:self syntenic dotplot construction using SynMap in CoGe (Data S1I, J). The *COI1* tandem pair appeared under GOs such as ‘response to external biotic stimulus’ (GO:0043207; Bonferroni p < 2.67e-08) and ‘defense response to other organism’ (GO:0098542; Bonferroni p < 3.5E-07).

**Figure 4.**
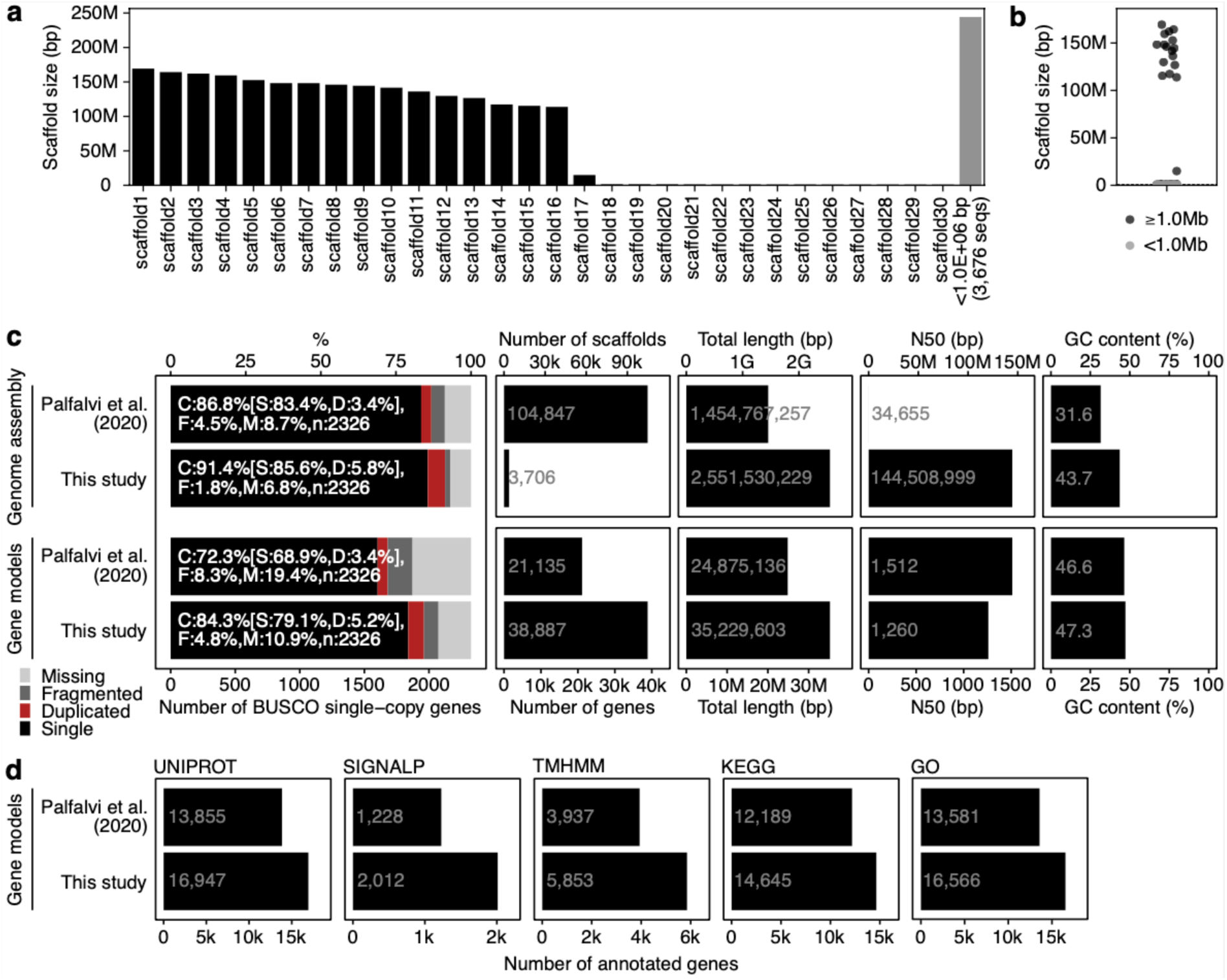
Chromosome-scale genome assembly of *Dionaea muscipula*. (a-b) Distribution of scaffold sizes. (c) Comparison of genome assembly and gene model statistics between the newly generated genome and a previously reported genome. (d) Comparison of annotated gene numbers using Trinotate. See also Figure S3 - S5 and Text S1.

Taking further advantage of this new assembly, we sequenced the DYSC^12^ mutant using Illumina short reads as well as two other *Dionaea* mutants, ‘Rose’ and ‘Basmati’, to characterize single-nucleotide variants (SNVs) and structural variants (SVs) potentially linked to their mutant phenotypes (Text S1).

Using this improved reference genome, we characterized differentially expressed genes (DEGs) in the unstimulated WT and DYSC mutant traps. While 864 genes were upregulated in DYSC compared to WT, 1071 genes were downregulated in this comparison (Figure 5 and Data S2A-E). According to MapMan annotation^18^, DEGs upregulated in DYSC mainly comprised genes associated with transcriptional regulation or solute transport. Interestingly, the most prominent groups of downregulated genes in DYSC were related to cell wall organization (bin 21) and enzyme classification (bin 50) (Figure 5, Data S2B-E). Among these, enzymes involved in biosynthesis of cell wall components were detected as well as cell wall modifying proteins such as expansins, which are known to induce creep and stress relaxation, thus conferring cell wall plasticity needed e.g. for cell expansion.

**Figure 5.**
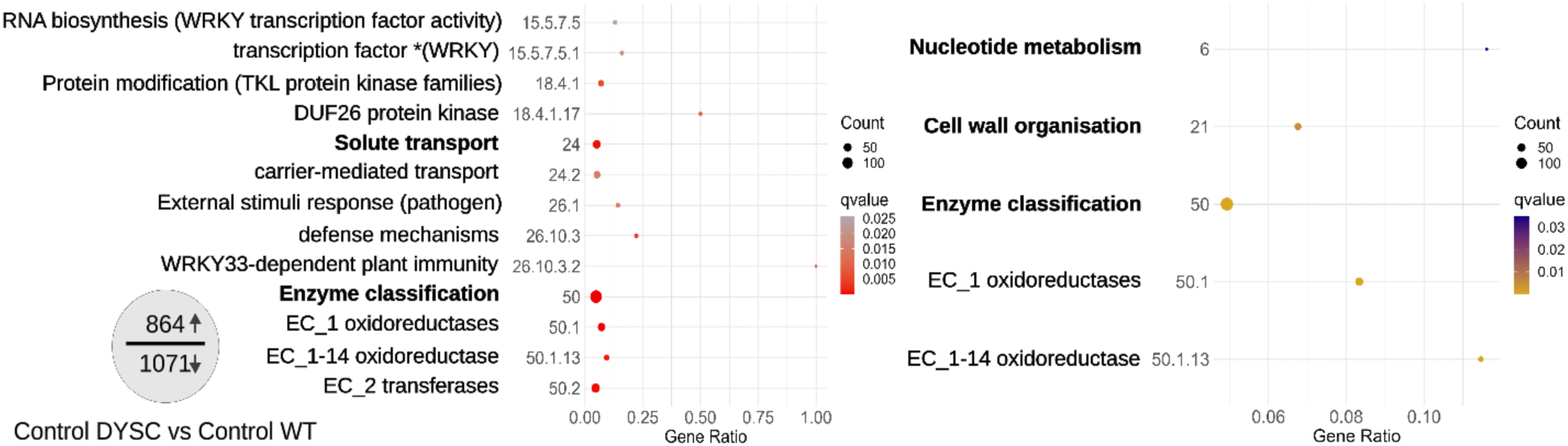
Differentially expressed genes and Mercator 4 Bin enrichment analysis of RNA-Seq Data. Enriched Mercator Bins of up- (left) and downregulated (right) genes in DYSC in the unstimulated ground state comparison. Enrichment based on Mercator 4 bin annotation of DEGs between DYSC and WT. Total number of DEGs is illustrated on the left. The y axis shows the different enriched Mercator Bins and the x axis the Gene Ratio. Gene Ratio refers to the ratio of enriched genes to all target genes. The number of enriched genes for each group is shown by dot size and color represents the qvalue. DEGs are defined as padj < 0.05 with log2FC > 1 and base Mean DYSC > 50 or log2Fc < −1 and base Mean WT > 50. Groups written in bold are supergroups. WT = *Dionaea muscipula* wildtype, DYSC = *Dionaea muscipula* DYSCALCULIA mutant/ cultivar, DEG = differentially expressed gene. See also Data S2.

### Altered cell wall properties in DYSC

DYSĆs altered trap characteristics may well impede proper trap deformation and thus snap buckling, since *Dionaea*’s trap closure mechanism is determined by leaf geometry^8,9^. In addition to this, fast trap closure is driven by simultaneous expansion of outer cell layers while inner cell layers are thought to collapse. Both processes require rapid changes in cell turgor and a certain cell wall plasticity, which allow distinct trap areas to change their geometry. We therefore wondered if mechanical cell wall properties might be altered in DYSC traps, thus preventing proper opening, i.e. pre-stress establishment, or the rapid conformational change underlying snap buckling. To analyze and compare mechanical cell wall properties of WT and DYSC traps, we applied cell indentation by AFM to determine Young’s moduli, a measure of cell wall elasticity, where a higher Young’s modulus corresponds to a stiffer (less elastic) cell wall.

AFM force-indentation curves were collected at three different positions, both on the inner and on the outer lobe surface (Figure 6, inlay, schematic trap): close to the midrib (position I), in the trap centre (position II) or at the leaf margins (position III). From the obtained force-indentation curves, Young’s moduli were then extracted using the Hertz model. In the trap centre (position II), there was no significant difference in elasticity between WT and DYSC for either side of the trap. In contrast, striking differences between WT and DYSC could be observed in the outer layer close to the midrib (position I) and in the inner layer close to the rim (position III). For these two specific positions, the Young’s Modulus for DYSC plants was significantly higher than that for WT plants at the same position. The schematic illustration of a *Dionaea muscipula* trap highlights the two distinct trap areas where the cell wall of DYSC mutants was significantly less elastic (i.e. stiffer) than for WT traps (Figure 6, inlay, schematic trap).

**Figure 6.**
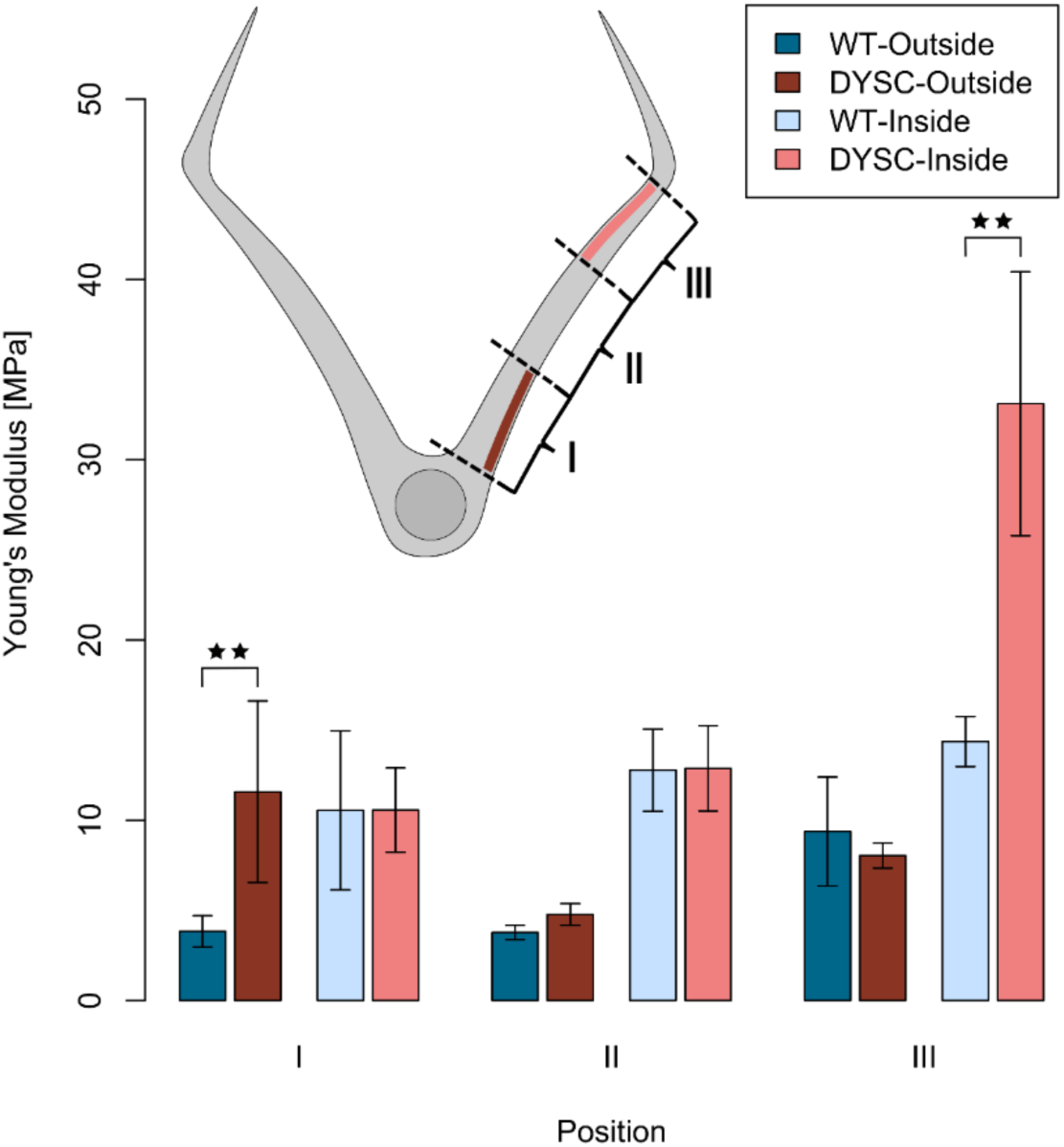
AFM reveals differences in the cell wall plasticity between DYSC and WT. AFM measurements of cell wall elasticity for WT and DYSC traps. Young’s Moduli were determined from AFM force-indentation curves at different trap positions (I, II, III, see schematic inlay). At each position, 10 force-indentation curves were recorded and the mean Young’s Modulus elasticity value from these 10 curves was calculated. Histogram bars represent the average (± standard deviation) from these mean Young’s moduli from n investigated trap lobes, n = 10-12 for WT and DYSC, for each of the three positions. WT (blue) and DYSC (red) traps were tested on the outer (darker colors) or inner face (lighter colors). Upper left: schematic cross section of a trap illustrating the different measuring positions. Position I is close to the midrib around the lowest trigger hair. Position II is located in the trap centre between the two upper trigger hairs. Position III marks the region below the rim of the trap. One freshly prepared trap was used for measurements at the three different positions, where the order in which the positions were probed was varied to exclude systematic error. Shapiro-Wilk p > 0.05; Mann-Whitney-test compares DYSC samples (red) against WT samples (blue) position dependent and surface dependent and is represented by stars: * p < 0.05, ** < 0.01, *** p < 0.001. WT = *Dionaea muscipula* wildtype, DYSC = *Dionaea muscipula* DYSCALCULIA mutant/ cultivar.

## Discussion

The Venus flytrap is known for one of the fastest movements in the plant kingdom in its carnivorous snap-traps: Mechanostimulation of touch-sensitive trigger hairs located on the inner trap surface by potential prey initiates both a Ca^2+^-wave and an action potential, simultaneously spreading all over the trap surface^1,2,6^. Decoding of both signals results in rapid trap closure by snap buckling, thus enabling effective prey capture. Quite recently, both experimental and simulation analyses revealed that the change in *Dionaea*’s trap conformation is very likely a result of simultaneous expansion of the outer epidermis and shrinkage of the inner epidermal cell layer ^9^. The establishment of a certain pre-stress state, i.e., an internal hydraulic pressure differential between these two layers, seems to be a prerequisite for proper snap buckling of the trap lobes. Interestingly, building up this pre-stress is obviously handicapped under conditions of reduced water supply in WT plants, and thus lower turgor pressure, which in turn results in impaired trap closure with respect to closure speed or geometrically correct lobe folding^9^.

In DYSC mutant plants, however, rapid trap closure by snap buckling is massively impaired even under ideal growth conditions, including good hydration. In contrast to what was described before, we found that DYSC traps can close upon mechanostimulation of trigger hairs, but closure takes much longer than in WT traps and is usually characterized by the absence of snap buckling. This defect does not depend on the nature of the applied stimulus, so that one might speculate that DYSC traps are generally unable to undergo rapid conformational change and thus snap closure.

In search for the mutation underlying DYSĆs phenotype, we scanned an updated WT reference genome and reanalyzed the DYSC mutant dataset for structural variations and single nucleotide variants. Compared to the genome version published earlier^23^, the current chromosome-scale assembly features more than 17,000 additional gene models, which should enhance genomic resolution of this and future studies on carnivorous plants. However, although we detected numerous SVs and SNVs, we were not able to attribute DYSĆs inability to snap close its trap to a single genetic mutation.

To bridge this gap, we analyzed our transcriptomic data using the improved version of the *Dionaea* genome as a reference. Comparing unstimulated DYSC to WT traps, more than 1900 genes were characterized as DEGs. Interestingly, many genes associated to cell wall biosynthesis or modification were identified among the genes downregulated in DYSC, such as expansins (Diomu_026145-T1, Diomu_034454-T1, Diomu_034452-T1) or proteins potentially involved in hemicellulose modification or -biosynthesis (Diomu_007193-T1, Diomu_031293-T1, Diomu_031149-T1). Targeting various cell wall components, expansins are well known to induce rapid wall loosening at low pH by disrupting bonds between laterally aligned cell wall polysaccharides ^47,48^. More than a decade ago it was already suggested that expansin-driven cell wall loosening might be involved in turgor-driven snap closure in *Dionaea*^49^. As such, DYSĆs incapacity to close by snap buckling might be affected by deregulation of expansins, i.e., via a mutation in a master regulator of the pathway(s).

Another way to modify cell wall plasticity is via methylesterification and -deesterification of pectins, which changes the free negative charge of homogalacturonan molecules and thus the dimension of crosslinking of pectin chains by Ca^2+^ ions. Therefore, the degree of homogalacturonan methylation affects the stiffness of the pectic matrix and thus cell wall elasticity. In numerous plants, pectin methylesterases (PME) and PME inhibitors (PMEI) have been characterized and described to be involved in cell wall modification during various processes ranging from growth processes to plant-environment interactions^50^. Interestingly, several PME/PMEIs are transcriptionally up- or downregulated in DYSC traps (Diomu_028350-T1, Diomu_028352-T1, Diomu_028356-T1, Diomu_025822-T1, Diomu_002489-T1). Therefore, *Dionaea*’s trap might feature different cell wall properties than WT traps, which may finally affect the mutant’s ability to rapidly undergo conformational trap changes associated with snap buckling, although mechanoperception and electrical signaling are not impaired in DYSC. However, detection of differentially expressed cell wall related genes in the whole trap does not provide the resolution required to explain potential differences in cell wall elasticity in distinct trap zones.

To investigate potentially different mechanical properties of DYSC and WT traps that may underlie the dysfunctional dynamic response to stimuli in DYSC traps, we applied AFM cell indentation. These studies provided information on the plasticity of WT and DYSC cell walls in different trap regions. While cell wall stiffness of WT and DYSC was similar in large parts of the trap, we detected significant alterations (i.e. enhanced cell wall stiffening) in DYSC traps in the outer layer close to the midrib and in the inner layer close to the rim. Taking into consideration that these are the trap zones undergoing pronounced conformational change during snap closure, i.e. differential shrinking and expansion in opposing inner and outer layers, enhanced stiffness in these regions may well impede snap buckling in DYSC. Since DYSC traps exhibit a smaller opening angle and reduced trap curvature, we cannot completely rule out, however, that pre-stress in DYSC already differs from WT traps. As mentioned earlier, proper pre-stress establishment is required to keep the trap in a “ready to snap”-mode^9^, so that DYSC traps might already be impaired by insufficient tension in the unstimulated ground state.

Considering DYSĆs impairment in Ca^2+^ decoding described earlier, its altered cell wall properties might be linked to Ca^2+^ signaling as well. Enzymes involved in cell wall remodeling such as PMEs/PMEIs, peroxidases or synthases of cell wall components might be transcriptionally or posttranslationally regulated by Ca^2+^ ions. Although PMEs/PMEIs as well as other cell wall modifying proteins such as expansins or Xyloglucan endotransglycosylases (e.g. TCH4) have not been described to directly bind calcium, calcium-dependent protein kinases (CDPKs) might regulate their activity, thus affecting cell wall plasticity required for proper snap buckling and trap function. Future studies on cell wall proteins under control of (stress) Ca^2+^ signalling are needed to reveal the platforms interconnecting mechanical and Ca^2+^ signalling.

Alternatively, proteins involved in rapid turgor changes required for the conformational change during snap buckling might be dysfunctional in DYSC: we cannot rule out that ion fluxes and turgor changes associated with trap buckling are regulated by Ca^2+^, so that this signalling cascade may be affected in DYSC as well. This hypothesis will become testable in the very near future using emerging high resolution turgor monitoring technology^51^.

## Acknowledgments

We acknowledge the following sources for funding: the Sofja Kovalevskaja programme of the Alexander von Humboldt Foundation (to K.F.), a Human Frontier Science Program (HFSP) Young Investigators Grant RGY0082/2021 (to K.F.), JSPS KAKENHI 23K20050 (to K.F.), a DFG Koselleck grant no. 415282803 (to R.H.). VAA acknowledges award 2030871 from the United States National Science Foundation. We thank Brigitte Neumann, Sarah Zehnter, Yoshie Kishida, and Akiko Watanabe for technical assistance. Computations were partially performed on the National Institute of Genetics (NIG) supercomputer, the Data Integration and Analysis Facility at National Institute for Basic Biology, the Erlangen National High Performance Computing Center, and High-Performance Computing Clusters at the University of Würzburg.

## Author Contributions

## Data availability

The chromosome-scale *Dionaea* genome assembly and gene models are available from the DNA Data Bank of Japan (DDBJ) with the accession numbers BAAGIJ010000001 to BAAGIJ010003706. DNA sequencing reads were deposited to DDBJ (Data S1B).

