## Supplemental text, figures and methods for "Impaired trap closure in the counting-deficient Venus flytrap mutant DYSCALCULIA is caused by cell wall biomechanics"

### Supplement Trebing et al.

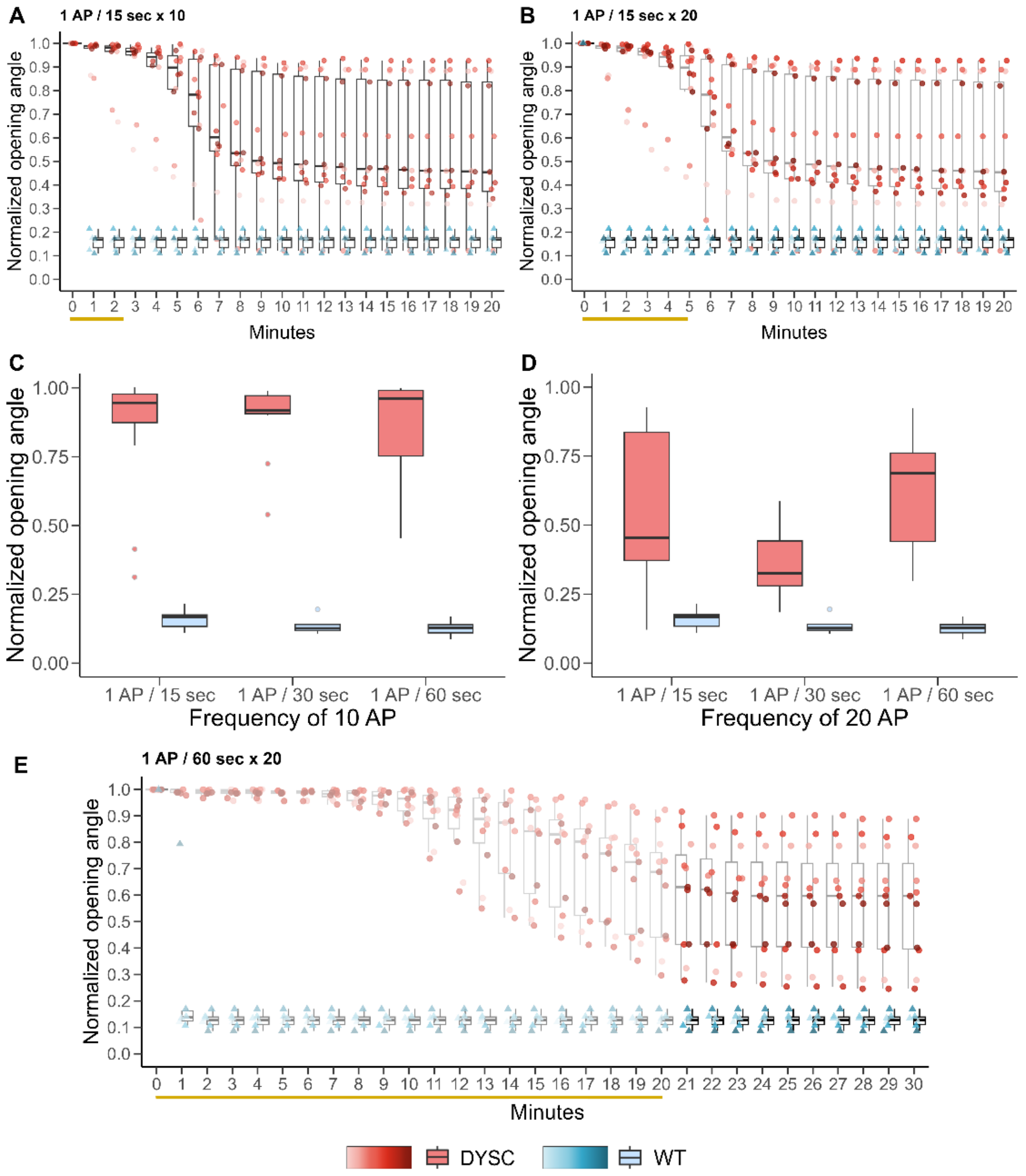

**Figure S1. Prolonged mechanostimulation activates trap closure in DYSC traps.**
Normalized opening angle after mechanostimulation of traps with a frequency of 1 AP / 15 sec for (A) 10 AP or (B) 20 AP for WT (blue) and DYSC (red). (C) and (D) show the final opening angles derived from Fig. 1 and S1A and S1B in response to different frequencies or AP numbers (C, 10 AP; D, 20 AP) as boxplots. Each trap´s opening angle was normalized to its initial opening angle. For normalization, 1 = completely open trap and 0 = sealed trap. Boxplots show the distribution of samples 20 min after starting mechanostimulation. (E) indicates the response to 20 APs at a frequency of 1 AP / 60 sec for up to 30 min. Highlighted data is a continuation of the data shown in Figure 1D up to 30 minutes. Opening angles of all traps are normalized to their initial opening angle. For normalization, 1 = completely open trap and 0 = sealed trap. 2 APs in WT samples led to fast trap closure in all cases. Orange bars below the x-axis indicate duration of mechanostimulation for DYSC samples. The transparent section is also shown in Figure 1D. Boxplots include the median (horizontal line), the upper / lower quartiles and whiskers. Colored dots show the outliers per analyzed group. n = 12 for DYSC and n = 6 for WT for each group. WT = *Dionaea muscipula* wildtype. DYSC = *Dionaea muscipula* DYSCALCULIA mutant/ cultivar. Related to Figures 1 and S2.

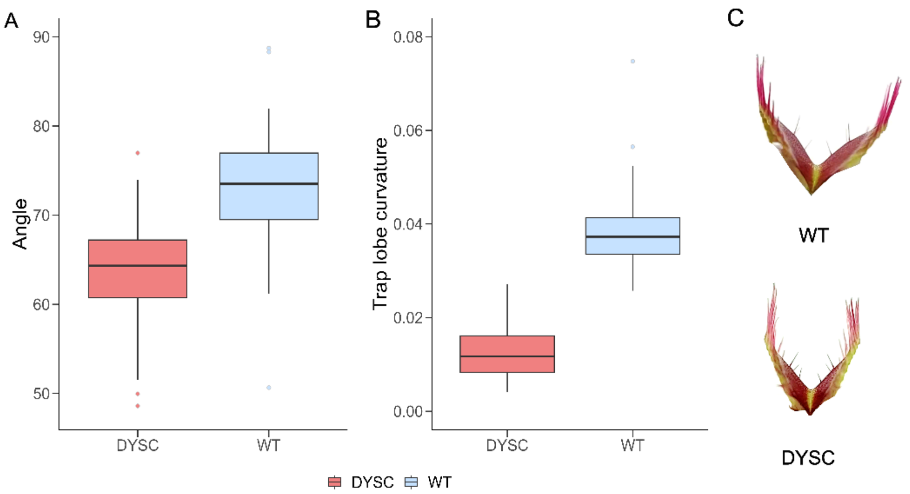

**Figure S2**. **Different starting parameters for DYSC and WT in the unstimulated ground state.**
Opening angle (A) and trap lobe curvature (B) in unstimulated, ground state WT (blue) and DYSC (red) traps. Boxplots include the median (horizontal line), the upper / lower quartiles and whiskers. Colored dots show the outliers per analyzed group. Lobe curvature was analyzed for both lobes of each trap, measured with the Kappa ImageJ plugin and represented by the average curvature ($mm^{-1}$). In C, representative traps are shown to illustrate the fundamental difference in trap geometry in both cultivars at the unstimulated resting state. For A n = 35-70 and for B n= 36-54. WT = *Dionaea muscipula* wildtype. DYSC = *Dionaea muscipula* DYSCALCULIA mutant/ cultivar. Related to Figure 3.

**Video S1**. **Exemplary sample setup of WT and DYSC traps to observe closure duration when placing traps into 100% EtOH.**
Closure duration was defined as the interval from first movement of trap lobes until the spines at the trap margin crossed. Video speed is 30 fps and is related to Figure 3 and Video S2.

**Video S2. Different trap conformational changes observed between WT and DYSC traps.**
Differences were found in experimental setup for EtOH Experiment and is related to Figure 3 and Video S1.

A

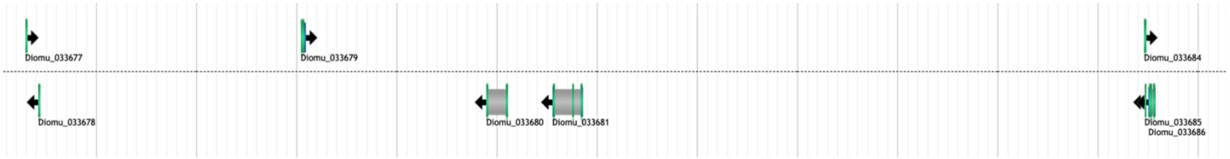

B

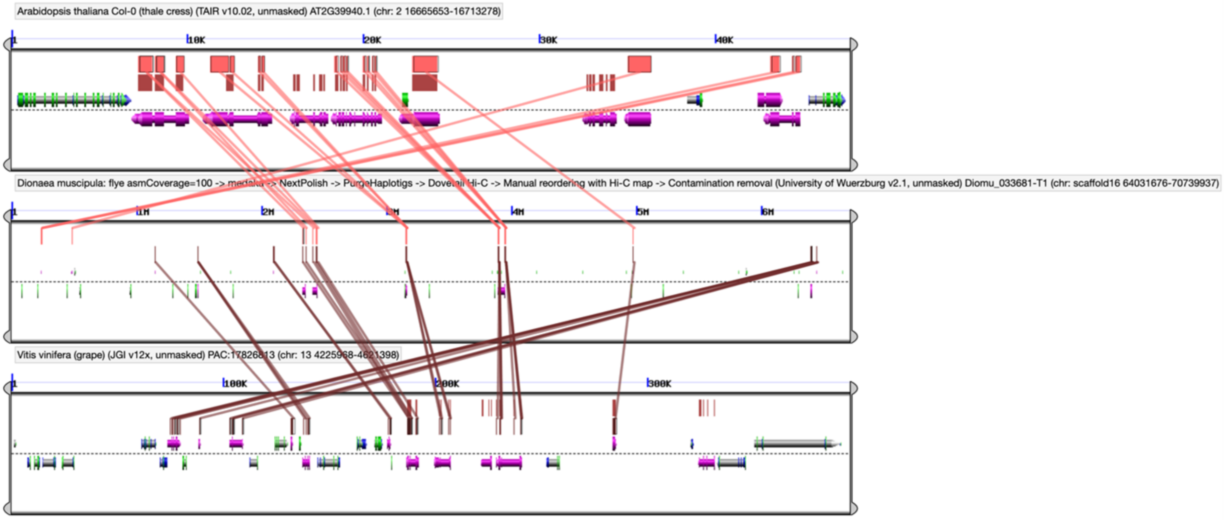

**Figure S3**. **Tandem duplication of COI1 homologs in *Dionaea muscipula*.**

A), genome browser view of a portion of scaffold16 of the *Dionaea* reference genome. Near the center lies a tandemly duplicated pair of *COI1* homologs. Diomu_033681 is likely a complete gene model, while Diomu_033680 appears to be truncated. Diomu_033680 may have been predicted in error or may be a pseudogene or otherwise truncated derivative with a deleted intron and exon. B) A CoGe GEvo microsynteny view of the same region (center) against homologous syntenic blocks in Arabidopsis (top) and grapevine, *Vitis vinifera* (lower block). Unlike *Dionaea*, *Arabidopsis* and *Vitis* possess only single *COI1* homologs in this view. Purple gene model coloration indicates genes joined by blast HSPs. Red HSPs and syntenic lines join *Arabidopsis* and *Dionaea* syntenic homologs, while brown HSPs and lines connect *Dionaea* and *Vitis* syntenic homologs. In the *Dionaea* block, center, the *COI1* tandem pair lies close to the right of the 2 Mb mark. Related to Figure 4.

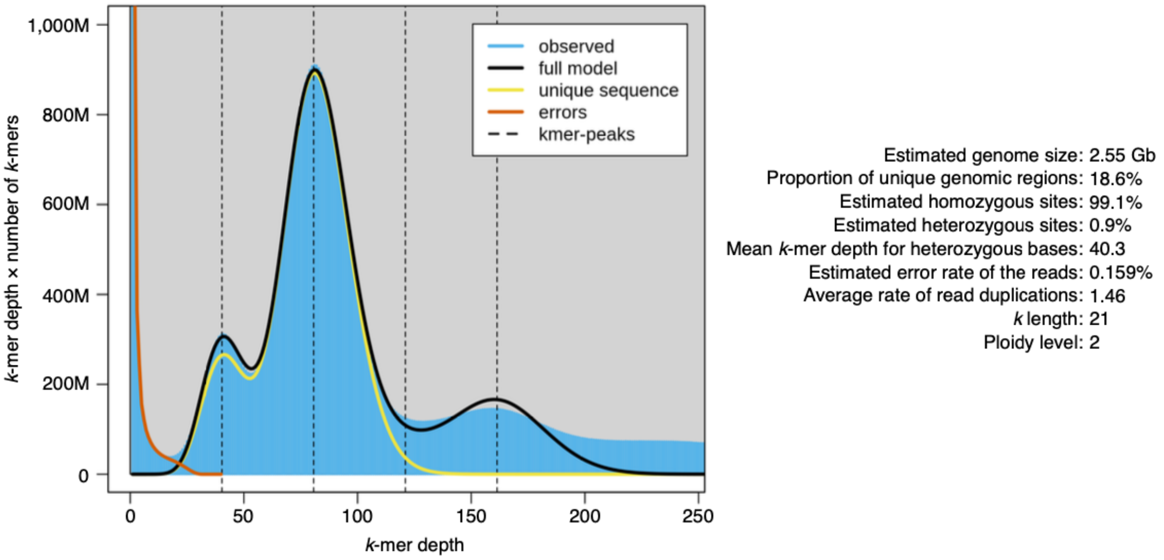

**Figure S4. The k-mer frequency distribution of the *Dionaea* genome.**
Illumina short reads were analyzed using GenomeScope v2.0. Related to Figure 4, Text S1.

**Text S1.** Structural and single nucleotide variant detection.
With a high-quality, chromosome-scale genome assembly available, we explored the potential role of highly disruptive mutations as an underlying cause or causes of the DYSC phenotype. This approach included analyses of structural variants (SVs) and single nucleotide variants (SNVs). Additionally, we sequenced the genomes of two *Dionaea muscipula* cultivars, 'Basmati' and 'Rose,' both of which lack critical trap structures. *Dionaea muscipula* 'Basmati', notably, does not feature trigger hairs, while *Dionaea muscipula* 'Rose' fails to develop the characteristic bi-lobed traps, instead forming leaves resembling non-carnivorous vegetative foliage, sometimes with serrated or toothed edges. These cultivars may therefore provide valuable insights into the genetic basis of trap development.
We identified a total of 102 SVs across the DYSC genome, 85 of which were classified as deletions and 17 as insertions (Data S1C). While most SVs were partially affecting coding regions, with variant lengths in the low hundreds bp, we identified several massive SVs. The largest SV we found was a 199-kbp deletion on scaffold6, effectively purging 5 genes in their entirety from the DYSC mutant. Among those were Diomu_012747 with high similarity to the general regulatory factor (GRF) gene family, two tetratricopeptide-repeat thioredoxin-likes (Diomu_012748, Diomu_012749), as well as two genes without detectable homology to *Arabidopsis thaliana* genes (Diomu_012746, Diomu_012750).
In the reference WT genome, these *GRF* and *TTL* genes are indeed present, and notably, as a segmentally duplicated gene set separated by ~200 kbp (Figure S5). In this block, the GRF homolog (Diomu_012747) and the *TTL*-like pair (Diomu_012748 and Diomu_012749) are tandemly repeated as Diomu_012743 and Diomu_012744, the latter of which is probably an erroneous fusion in the annotation of two *TTL*-like gene models. The TTL homologs could be of importance to the impaired cell wall plasticity phenotype of the DYSC mutant, as members of this family, such as *TTL1* have been reported to be implicated in cell wall elasticity in Arabidopsis, although this has only been demonstrated for roots[41]. Interestingly, scaffold 6 is host to another massive, 165-kbp large deletion in DYSC, although the only deleted gene, a *SEC7*-like guanine nucleotide exchange family protein (Diomu_013375), is unlikely to be related to the DYSC phenotype. Strangely, there appears to be detectable expression for the genes discussed above, in particular of the *GRF* homolog (Diomu_012747), but we cannot exclude the possibility that these represent RNA-Seq reads from the segmentally duplicated paralogs in the homologous gene cluster nearby. Additionally, while Diomu_012747 overlaps a predicted deletion, coverage in the area is

low in all samples, and SV inference at this site remains uncertain (Figure S5C). If we restrict our search by only taking trace or non-expression in the DYSC mutant into account (TPM < 0.5), the list of candidates shrinks down to 22 genes. Most notably among these are Diomu_011798 (scaffold5) and Diomu_020163 (scaffold9), bearing high sequence similarity to calcium dependent protein kinases (*CPK*) and a chitin responsive *PHD* finger transcription factor (*AT5G58610*) respectively, believed to function in plant defense[42].

Since SNVs encompass all variations of single nucleotides between genotypes, regardless of their impact on a coding region of the genome, they may inflate the dataset with numerous hits not seemingly relevant to the DYSC phenotype. Hence, we focused our search on genes that acquired a premature stop codon, effectively cutting a gene product short and making it defunct. In total, we found 87 genes containing a high-confidence premature stop-codon SNV (Data S1F). While the affected genes covered a wide variety of functions, no single candidate provided a satisfying explanation for the DYSC phenotype.

We identified 192 SVs (128 deletions and 64 insertions) in the *Dionaea muscipula* ‘Rose’ genome, with sizes ranging from 50 bp to 18 kbp (Data S1D). Several SVs affected genes involved in meristem organization and differentiation, including genes belonging to the small *ULTRAPETALA* (*ULT*) gene family (Diomu_019791), a p60 katanin protein (Diomu_004364), and a *BREAST CANCER ASSOCIATED RING* (*BARD*) gene (Diomu_032438). *KATANIN/ERH3 (Ectopic Root Hair 3)* is well characterized in *Arabidopsis thaliana* for its role in microtubule array regulation and is critical for the development of multiple plant structures, including embryos, roots, meristems, and leaves[43]. *ULT1* functions in a context-dependent manner, influencing meristem size and floral development via interaction with *WUSCHEL*. *BARD1* regulates *WUSCHEL* expression and is crucial for shoot apical meristem (SAM) organization[44]. The same study demonstrated that *BARD1* is essential for adaxial–abaxial leaf polarity, implying that disrupted meristem function may underlie the malformed leaves in *Dionaea muscipula* ‘Rose’.

Additionally, we identified 268 genes containing premature stop codons (Data S1G). Among them was a putative KNOX gene (Diomu_027934) with similarity to *BREVIPEDICELLUS* (*BP*, also known as *KNOTTED-LIKE FROM ARABIDOPSIS THALIANA KNAT1*) a gene involved in leaf development and shape regulation via interactions with auxin and the *ASYMMETRIC LEAVES1* transcription factor[45]. Overexpression of *BP* in *Arabidopsis thaliana* leads to lobed leaves, making it a potential candidate for contributing to the aberrant leaf morphology of *Dionaea muscipula* ‘Rose’.

In contrast to ‘Rose’ or DYSC, relatively few SVs were detected in *Dionaea muscipula* ‘Basmati’ (26 total: 20 deletions and 6 insertions) (Data S1E). However, SNV analysis revealed an unexpectedly high number of premature stop codons, affecting 1,109 genes (Data S1H). Given that trigger hairs in *Dionaea* are thought to be functionally analogous to the glandular tentacles of *Drosera*, we searched for genes implicated in trichome development. We identified a gene belonging to the *INDOLE-3-ACETIC ACID INDUCIBLE* (IAA) family (Diomu_005068). Members of this family, such as IAA15 were shown to regulate glandular trichome density in *Solanum lycopersicum*[46]. Since trichome development is induced by gibberellic acid (GA), disruptions in GA signaling could explain the absence of trigger hairs in ‘Basmati’. Supporting this hypothesis, we identified a putative GA-receptor (Diomu_012234). However, whether these genes account for the phenotype remains speculative, especially since neither anatomical nor molecular homology has been established between *Solanaceae* or *Arabidopsis* trichomes and either *Drosera* or *Dionaea* tentacles or glands. Given the high number of SNVs, it may be that multiple gene disruptions collectively contribute to the loss of trigger hairs in ‘Basmati’.

Related to Figure 4 and Figure S4.

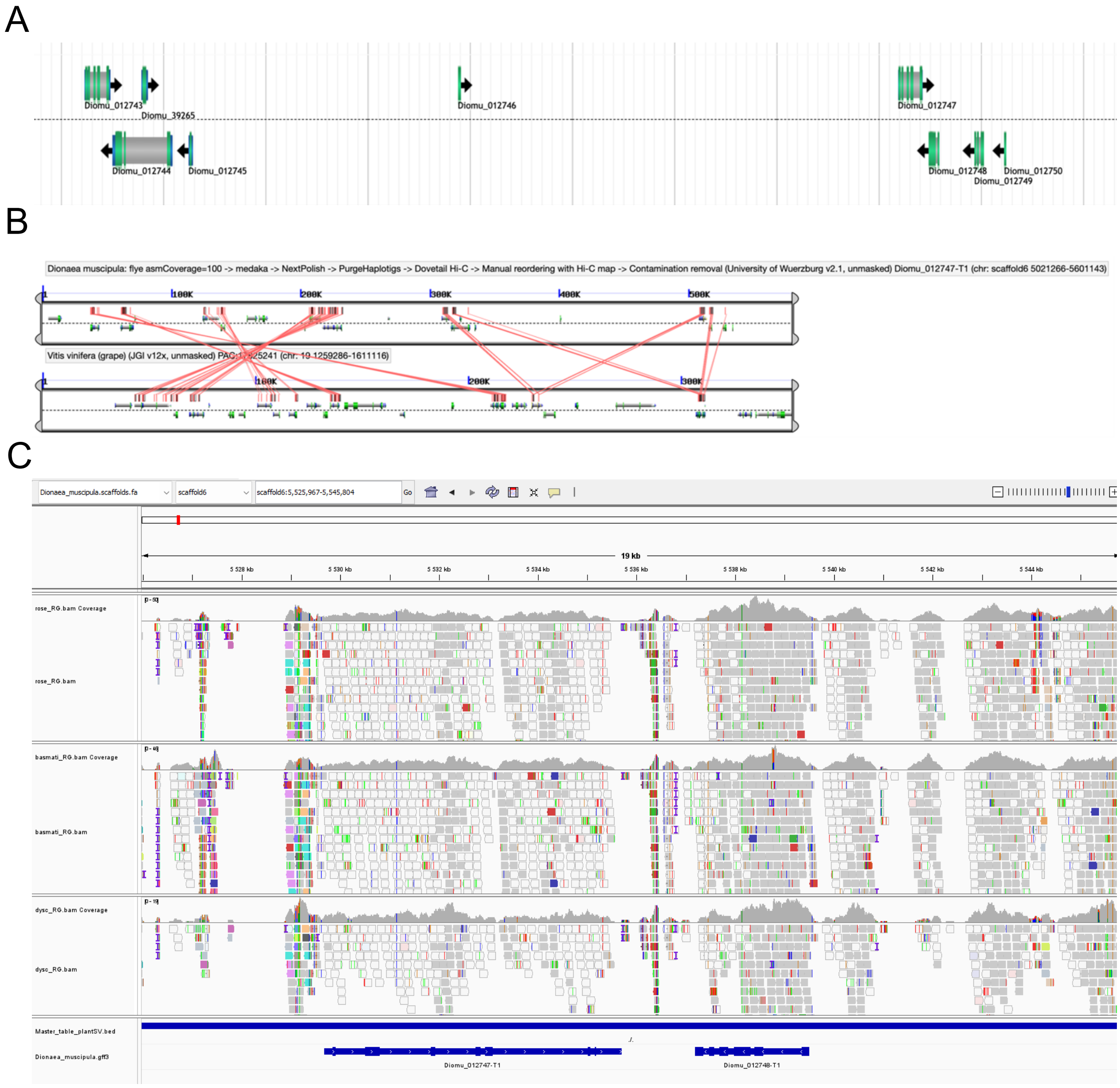

**Figure S5. Segmental duplication and putative deletion of GRF/TTL gene clusters in *Dionaea muscipula*.**

A) Genome browser view of a portion of scaffold6 of the *Dionaea* reference genome. To the right lie the *GRF* homolog (Diomu_012747) and *TTL*-like gene pair (Diomu_012748 and Diomu_012749) that are deleted in the DYSC mutant. Notably, however, these genes are segmentally duplicated together to the left (Diomu_012743 is a *GRF* paralog, and Diomu_012744 is likely an erroneous gene model representing a fused *TTL*-like paralog pair). B) A CoGe GEvo microsynteny view of the same region (top) against the homologous syntenic block in grapevine, *Vitis vinifera* (lower block). Blast HSPs are colored red as are syntenic lines connecting them. The segmentally duplicated *GRF/TTL* gene clusters lie close to the 300 Kb and 500 Kb marks in the *Dionaea* block, upper. The duplicated *Dionaea GRF/TTL* clusters have single matches in the *Vitis* genome, and these single *Vitis GRF* and *TTL* genes, while close to either, are separated by over 100 Kb of DNA and 3 intervening genes. Therefore, both the *GRF-TTL* gene clustering and their segmental duplication nearby are likely specific to the *Dionaea* lineage. C) Genome browser view of genomic region encompassing the Diomu_012747 locus and surrounding area flagged as deleted in our SNV analysis. Inspection of the region reveals mapped sequencing reads across the putatively deleted gene. While read presence suggests possible expression, overall read coverage is sparse, and base quality acr *Segmental duplication and putative deletion of GRF-TTL gene clusters in Dionaea muscipula.*oss this region is low. These factors complicate definitive interpretation of the deletion status. Related to Figure 4 and Text S1.

**Data S1, Transcriptomic resources used for gene modelling , Whole Genome Sequencing (WGS) experiment information and downstream SV and SNV analysis results for different *Dionaea muscipula* genotypes.** Related to STAR METHODS and Text S1

A. Sequence Read Archive (SRA) metadata for *Dionaea muscipula* RNA-Seq samples used in gene model prediction. All samples were taken from the publicly available bioproject PRJNA530242 and included trap tissue before and after feeding, as well as isolated trigger hairs and trap tissue with trigger hairs removed.
B. Bioproject and library information on *Dionaea muscipula* DNA-Seq data generated in this study, including *Dionaea muscipula* wildtype WGS and Hi-C experiments, as well as WGS experiments for the *Dionaea muscipula* DYSC mutant, *Dionaea muscipula* 'Rose' and *Dionaea muscipula* 'Basmati'.
C-E. Lists of genes predicted to contain structural variants (insertions or deletions) in the *Dionaea muscipula* DYSC (C) mutant, *Dionaea muscipula* 'Rose' (D) and *Dionaea muscipula* 'Basmati' (E) according to our SV analysis. Tables contain information on the SV type, size, location relative to the scaffold, as well as the ID and location of the affected genes.
F-H. Lists of genes predicted to contain pre-mature stop codons in the *Dionaea muscipula* DYSC (F) mutant, *Dionaea muscipula* 'Rose' (G) and *Dionaea muscipula* 'Basmati' (H) according to our SNV analysis. Annotated SNVs needed to satisfy a minimum quality score of 30, homozygosity, and a depth between 3 and 2× the average read coverage. Additionally, SNVs were filtered by predicted effect (i.e. introduction of premature stop-codons) and uniqueness towards the respective genotype. Tables include information on SNV location relative to the scaffold, the predicted nucleotide variant and the ID of the affected gene.
I. List of significantly enriched GO-terms in GO-term enrichment of *Dionaea muscipula* tandem duplicates ordered by Bonferroni corrected p-value.
J.Genes associated with the top 58 significantly enriched GO terms identified in Table I. This table lists all *Dionaea muscipula* genes contributing to the top 58 GO terms enriched in the analysis shown in Table I. For each gene, the corresponding best reciprocal BLAST hit in *Arabidopsis thaliana* is provided, along with its functional annotation.

**Video S2. RNA-Seq Data analysis of differentially expressed genes and Bin enrichment analysis of up- and downregulated DEGs for the comparison control DYSC vs control WT.** Related to Figure 5

**Data S2. RNA-Seq Data analysis of differentially expressed genes and Bin enrichment analysis of up- and downregulated DEGs for the comparison control DYSC vs control WT.** Related to Figure 5
A. Differentially expressed genes for control DYSC vs control WT. Definition DEG: padj < 0.05 with $\log_2$FC > 1 and base Mean DYSC > 50 for upregulated genes or padj < 0.05, $\log_2$Fc < -1 and base Mean WT > 50 for downregulated genes.
B. MapMan Bin enrichment of downregulated genes. All downregulated DEGs of the control comparison were submitted to https://www.plabipd.de/mercator_main.html. Two sided Fishers Exact test with an FDR-adjusted-p-value cutoff of 0.05 was set. Shown are all enriched groups of Mercator 4 for the downregulated genes list.
C. Downregulated genes of the enriched bins shown in Data S2B. Identifiers of each enrichment group were combined with general information to provide a better description of the individual downregulated genes.
D. MapMan Bin enrichment of upregulated genes. All upregulated DEGs of the control comparison were submitted to https://www.plabipd.de/mercator_main.html. Two sided Fishers Exact test with an FDR-adjusted-p-value cutoff of 0.05 was set. Shown are all enriched groups of Mercator 4 for the upregulated genes list.
E. Upregulated genes of the enriched bins shown in Data S2D. Identifiers of each enrichment group were combined with general information to provide a better description of the individual upregulated genes.

### STAR Methods

#### Key resources table

| REAGENT or RESOURCE | SOURCE | IDENTIFIER |
|---|---|---|
| Chemicals, peptides, and recombinant proteins | | |
| Coronatine | Sigma-Aldrich, Merck | Cat#C8115 |
| Ethanol | Carl Roth, Karlsruhe, Germany | Cat#P076.1 |
| Tris-HCl | PanReacAppliChem, Darmstadt, Germany | Cat#A1086 |
| EDTA-2Na | PanReacAppliChem, Darmstadt, Germany | Cat#131669 |
| KCl | PanReacAppliChem, Darmstadt, Germany | Cat#131494 |
| NaCl | PanReacAppliChem, Darmstadt, Germany | Cat#131659 |
| PVP-10 | SIGMA-Aldrich / Merck, Darmstadt, Germany | Cat#PVP10 |
| Spermine tetrahydrochloride | SIGMA-Aldrich / Merck, Darmstadt, Germany | Cat#85610 |
| Spermidine trihydrochloride | PanReacAppliChem, Darmstadt, Germany | Cat#A0673 |
| β -mercaptoethanol | SIGMA-Aldrich / Merck, Darmstadt, Germany | Cat#M3148 |
| Triton X-100 | SIGMA-Aldrich / Merck, Darmstadt, Germany | Cat#T8787 |
| NucleoBond HMW DNA Kit | Macherey-Nagel, Düren, Germany | Cat#740160.20 |
| Qubit dsDNA BR Assay kit | ThermoFisher Scientific | Cat#Q32850 |
| NEBNext Ultra II DNA Library Prep Kit | New England Biolabs, Frankfurt am Main,Germany | Cat#E7645 |
| Deposited data | | |
| RNA-Seq experiment | [1] | SRA: PRJNA899051 |
| Nanopore DNA-Seq WT samples | This Paper | SRA: PRJDB15741 |
| DNA-Seq DYSC, Rose, Basmati | This Paper | SRA: PRJDB15751 |
| RNA-Seq samples used for gene modelling | [2] | SRA: PRJNA530242 |
| Experimental models: Organisms/strains | | |
| *Dionaea muscipula* WT cultivar | Cresco Carnivora (Netherlands) | N/A |
| *Dionaea muscipula* DYSC cultivar | Green Jaws (Germany) | Cat# 104.000.065 |
| *Dionaea muscipula* Rose cultivar | Green Jaws (Germany) | Cat#104.000.088 |
| *Dionaea muscipula* Basmati cultivar | Green Jaws (Germany) | Cat#104.000.061 |
| Software and algorithms | | |
| Fiji / ImageJ2 (version 2.3.0/1.53f) | [3] | https://imagej.net/software/fiji/downloads |
| FastQC (version 0.12.1) | [4] | https://www.bioinformatics.babraham.ac.uk/ projects/fastqc/ |
| MultiQC (version 1.12) | [5] | https://github.com/MultiQC/MultiQC |

| | | |
|---|---|---|
| AMALGKIT (version 0.12.0) | Unpublished | https://github.com/kfuku52/amalgkit |
| RStudio (version 4.3.3) | [6] | https://cran.r-project.org/bin/windows/base/old/4.3.3/ |
| DESeq2 package (version 1.4.4) | [7] | https://github.com/mikelove/DESeq2 |
| Mercator 4 (version 6) | [8] | https://www.plabipd.de/mercator_main.html |
| Mercator 3.6 | [9] | https://www.plabipd.de/mercator_main.html |
| Flye (version 2.8.3) | [10] | https://github.com/mikolmogorov/Flye |
| GenomeScope (version 2.0) | [11] | https://github.com/schatzlab/genomescope |
| Medaka (version 1.4.3) | Oxford Nanopore Technologies Ltd, Oxford, United Kingdom | https://github.com/nanoporetech/medaka |
| NextPolish (version 1.3.1) | [12] | https://github.com/Nextomics/NextPolish |
| RepeatModeler (version 2.0.2) | [13] | https://github.com/Dfam-consortium/RepeatModeler |
| Purge Haplotigs (Version 1.1.2) | [14] | https://bitbucket.org/mroachawri/purge_haplotigs/src |
| MMseq2 (version 13.45111) | [15] | https://github.com/soedinglab/MMseqs2 |
| RepeatMasker (version 4.0.9) | [16] | https://github.com/Dfam-consortium/RepeatMasker |
| Funannotate (version 1.8.14) | [17] | https://github.com/nextgenusfs/funannotate |
| Trinity (version 2.8.5) | [18] | https://github.com/trinityrnaseq/trinityrnaseq/wiki |
| Augustus (version 3.3.3) | [19] | https://github.com/Gaius-Augustus/Augustus |
| GeneMark-ES (version 4.71) | [20] | https://genemark.bme.gatech.edu/gmes_instructions.html |
| Trinotate (version 3.2.1) | [21] | https://github.com/Trinotate/Trinotate |
| DIAMOND BLASTP search (version 2.1.9) | [22] | https://github.com/bbuchfink/diamond |
| BWA (version 0.7.17) | [23] | https://github.com/lh3/bwa |
| BCFtools (version 1.15.1) | [24] | https://github.com/samtools/bcftools |

| | | |
|---|---|---|
| SnpEff (version 5.1d) | 25 | https://github.com/pcingola/SnpEff |
| Manta (version 1.6) | 26 | https://github.com/Illumina/manta |
| BayesTyper (version 1.5) | 27 | https://github.com/bioinformatics-centre/BayesTyper |
| VCFtools (version 0.1.16) | 28 | https://github.com/vcftools/vcftools |
| CoGE | 29 | CoGe: Comparative Genomics |
| SynMap | 30 | CoGe: SynMap |
| GOATOOLS | 31 | https://github.com/tanghaibao/goatools |
| Other | | |
| Samsung Galaxy A71 | Samsung | Galaxy A71 |
| Fujifilm x-T2 camera | Fujifilm | X-T2 |
| Micro Star 30R | VWR | Cat#521-2574 |
| Molecular Force Probe 3D (MFP-3D) AFM | Asylum Research, Oxford Instruments | |
| SD-R150-NCL tip | Nanosensors | SD-R150-NCL |
| Nylon filter with 100 μm mesh size, Falcon Cell Strainers, | Corning | Cat#352360 |
| 2100 Bioanalyzer | Agilent | |

references (supplement)

1. Iosip AL*, et al.* DYSCALCULIA, a Venus flytrap mutant without the ability to count action potentials. *Curr Biol* **33**, 589-596 e585 (2023).

2. Procko C*, et al.* Stretch-activated ion channels identified in the touch-sensitive structures of carnivorous Droseraceae plants. *Elife* **10**, 10:e64250 (2021).

3. Rueden CT*, et al.* ImageJ2: ImageJ for the next generation of scientific image data. *BMC Bioinformatics* **18**, 529 (2017).

4. Andrews S. FastQC: a quality control tool for high throughput sequence data.). Babraham Bioinformatics, Babraham Institute, Cambridge, United Kingdom (2010).

5. Ewels P, Magnusson M, Lundin S, Käller M. MultiQC: summarize analysis results for multiple tools and samples in a single report. *Bioinformatics* **32**, 3047-3048 (2016).

6. Team RC. R: A language and environment for statistical computing. (2013).

17. Palmer JM, Stajich J. Funannotate v1.8.1: Eukaryotic genome annotation (v1.8.1). Zenodo., (2020).

23. Li H. Aligning sequence reads, clone sequences and assembly contigs with BWA-MEM. (2013).
